# SMELL-S and SMELL-R: olfactory tests not influenced by odor-specific insensitivity or prior olfactory experience

**DOI:** 10.1101/161000

**Authors:** Julien W. Hsieh, Andreas Keller, Michele Wong, Rong-San Jiang, Leslie B. Vosshall

## Abstract

Smell dysfunction is a common and underdiagnosed medical condition that can have serious consequences. It is also an early biomarker of Alzheimer’s disease that precedes detectable memory loss. Clinical tests that evaluate the sense of smell face two major challenges. First, human sensitivity to individual odorants varies significantly, leading to potential misdiagnosis of people with an otherwise normal sense of smell but insensitivity to the test odorant. Second, prior familiarity with odor stimuli can bias smell test performance. We have developed new non-semantic tests for olfactory sensitivity (SMELL-S) and olfactory resolution (SMELL-R) that overcome these challenges by using mixtures of odorants that have unfamiliar smells. The tests can be self-administered with minimal training and showed high test-retest reliability. Because SMELL-S uses odor mixtures rather than a single molecule, odor-specific insensitivity is averaged out. Indeed, SMELL-S accurately distinguished people with normal and dysfunctional smell. SMELL-R is a discrimination test in which the difference between two stimulus mixtures can be altered stepwise. This is an advance over current discrimination tests, which ask subjects to discriminate monomolecular odorants whose difference cannot be objectively calculated. SMELL-R showed significantly less bias in scores between North American and Taiwanese subjects than conventional semantically-based smell tests that need to be adapted and translated to different populations. We predict that SMELL-S and SMELL-R will be broadly effective in diagnosing smell dysfunction, including that associated with the earliest signs of memory loss in Alzheimer’s disease.

**Significance statement:** Currently available smell testing methods can misdiagnose subjects with lack of prior experience or insensitivity to the odorants used in the test. This introduces a source of bias into clinical tests aimed at detecting patients with olfactory dysfunction. We have developed smell tests that use mixtures of 30 molecules that average out the variability in sensitivity to individual molecules. Because these mixtures have unfamiliar odors, and the tests are non-semantic, their use eliminates differences in test performance due to the familiarity with the smells or the words used to describe them. The SMELL-S and SMELL-R tests facilitate smell testing of diverse populations, without the need to adapt the test stimuli.

## Introduction

Smell dysfunction manifests itself primarily in the reduced ability to detect or identify volatile chemicals, and ranges from the complete inability to smell any odors, to a partial reduction in olfactory sensitivity, to smell distortion, for instance that a large number of odors smell like cigarette smoke. The prevalence of smell dysfunction in the general adult population is about 20% in Europe and the United States (1–3). This condition is dangerous because those affected are unable to detect fire, spoiled food, hazardous chemicals, and leaks of odorized natural gas (4, 5). Smell loss also has severe health consequences, including mental health symptoms such as depression, anxiety, and social isolation. It affects quality of life by altering food preferences and the amount of food ingested (5). Food is often perceived as bland or tasteless by patients with smell disorders, leading to loss of appetite or overeating (4, 5).

Smell dysfunction has many causes, including head trauma, upper respiratory tract infection, nasal polyps, and congenital anomalies (6, 7). In many cases, the cause of smell dysfunction is unknown (5, 8). Importantly, smell dysfunction is an early sign of Alzheimer’s disease (9), the most common cause of dementia in the United States that is projected to affect an estimated 1 in every 45 individuals by 2050 (10). There is growing evidence that diminished olfactory function arises early in the progression of Alzheimer’s disease, and is highly predictive of future cognitive decline (3, 4). Because of the high prevalence and dramatic consequences of smell loss, accurate diagnosis of olfactory dysfunction is important. While self-reported hearing loss tends to be accurate (11), self-reporting of olfactory dysfunction is notoriously unreliable. Therefore, accurate diagnostic tests for smell dysfunction that can be deployed worldwide are critically important. Following a diagnosis, therapeutic options and counselling can be offered to patients suffering from smell loss (12).

In clinical smell testing, patients are presented with odor stimuli in a variety of formats, including scratch ‘n’ sniff strips, glass vials or jars, felt-tip pens, or paper scent strips used in perfume shops, and asked to answer questions about what they smell. Smell tests assess the ability of subjects to detect, discriminate, or identify odors. Olfactory threshold tests measure the lowest concentration of an odor stimulus that a patient can perceive, while discrimination tests assess the ability of subjects to distinguish two different smells. Finally, odor identification tests evaluate whether a patient can detect and match odors to standard words that describe the smell (13).

There are two major challenges to reliably testing a patient’s sense of smell. First, sensitivity to monomolecular odorants varies greatly even among subjects with a normal sense of smell (14- 16). All commercial smell tests that use monomolecular odorants therefore run the danger of misdiagnosing patients. For instance, when a patient has a low score on a test that assesses olfactory sensitivity with the rose-like odor phenyl ethyl alcohol (17), it is difficult to know whether the patient suffers from general smell dysfunction, or is merely insensitive to phenylethyl alcohol with an otherwise normal sense of smell.

The second challenge is to develop a test that is not influenced by the patient’s prior olfactory experiences. This has an obvious influence on the results of odor identification tests such as the University of Pennsylvania Smell Identification Test (UPSIT) for which subjects are given a booklet with 40 scratch ‘n’ sniff items and asked to select one of four words (for example “gingerbread”, “menthol”, “apple”, or “cheddar cheese”) that best describes what the odor smells like. Whether a patient can correctly identify the smell of gingerbread depends not only on the patient’s sense of smell, but also on whether the patient has previously encountered the smell of gingerbread. This in turn depends on many factors such as the cultural and age group the patient belongs to. To address this familiarity problem, the UPSIT has been adapted for use in a number of countries worldwide by replacing unfamiliar items and adapting the answers on the multiple-choice test. For instance, the North American UPSIT was adapted for Taiwanese subjects by replacing “clove”, “cheddar cheese”, “cinnamon”, “gingerbread”, “dill pickle”, “lime”, “wintergreen”, and “grass” with “sandalwood”, “fish”, “coffee”, “rubber tire”, “jasmine”, “grapefruit”, “magnolia”, and “baby powder” (18). The strong influence of prior olfactory experience on the test results limits the utility of odor identification tests. Even performance on non-semantic odor discrimination tasks depends on prior experience with the odorants (19, 20), and it is therefore important to avoid stimuli having differential familiarity in the test population.

We have developed two non-semantic smell tests that meet both challenges by using mixtures of odorous molecules that subjects perceive as unfamiliar. SMELL-S measures olfactory sensitivity, the ability to detect increasing dilution of the mixture of odorants. SMELL-R is an olfactory resolution test that measures the ability of subjects to discriminate the smell of two mixtures that become progressively more similar as the test proceeds. Neither of the tests requires that subjects match words with a smell percept. We show that SMELL-S and SMELL-R are highly reliable olfactory tests that overcome problems with odor-specific insensitivity, and that can be applied without adaptation to subjects in a different country. We expect that these tests, applied in combination, will be highly effective for early diagnosis of smell dysfunction in different populations.

## Results

### Designing two new smell tests: SMELL-S and SMELL-R

To improve currently available diagnostic tools for testing olfactory function, we created two new smell tests based on odorant mixtures. The Olfactory Sensitivity Test (SMELL-S) measures sensitivity to a mixture of 30 monomolecular odorants (Fig. 1*A*). The Olfactory Resolution Test (SMELL-R) measures the ability of subjects to discriminate the smell of such mixtures with an increase in overlapping components (Fig. 1*B*) (21, 22). Tests were presented in glass jars or vials as triangle tests, in which subjects were asked to pick out the stimulus with the strongest odor (SMELL-S) or the odd odor (SMELL-R). Both tests used adaptive staircase procedures that are standard in clinical olfactory testing (23) (Fig. 1*A*, 1*B*).

**Fig. 1.**
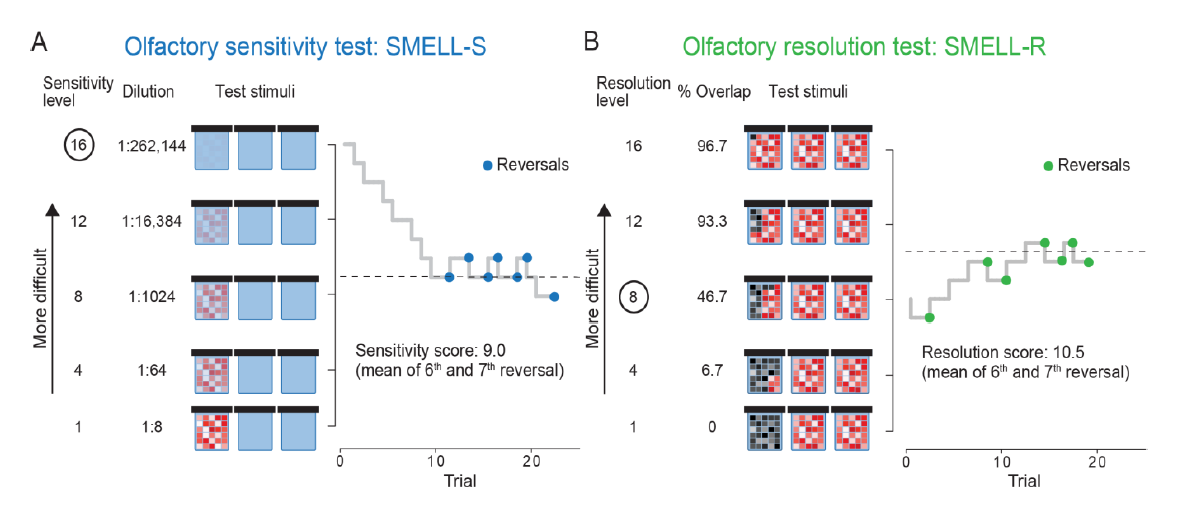
SMELL-S olfactory sensitivity and SMELL-R olfactory resolution tests. (*A*) Schematic of triangle test stimuli for SMELL-S, comprising two glass vials containing solvent (blue) and one containing increasingly diluted mixtures of 30 molecules (red-white mosaic). Olfactory sensitivity of a subject measured with SMELL-S [Subject Expt 1-A023, SMELL-S (v2)]. (*B*) Schematic of triangle test stimuli for SMELL-R, comprising two jars containing the same mixture of 30 molecules (red-white mosaic) and a one containing mixtures of 30 molecules (black-gray mosaic) with an increasing number of molecules shared with the other two. Olfactory resolution of a subject measured with SMELL-R [Subject Expt 1-A016, SMELL-R (v1)]. Circles in A and B indicate starting level for each test.

**Fig. 2.**
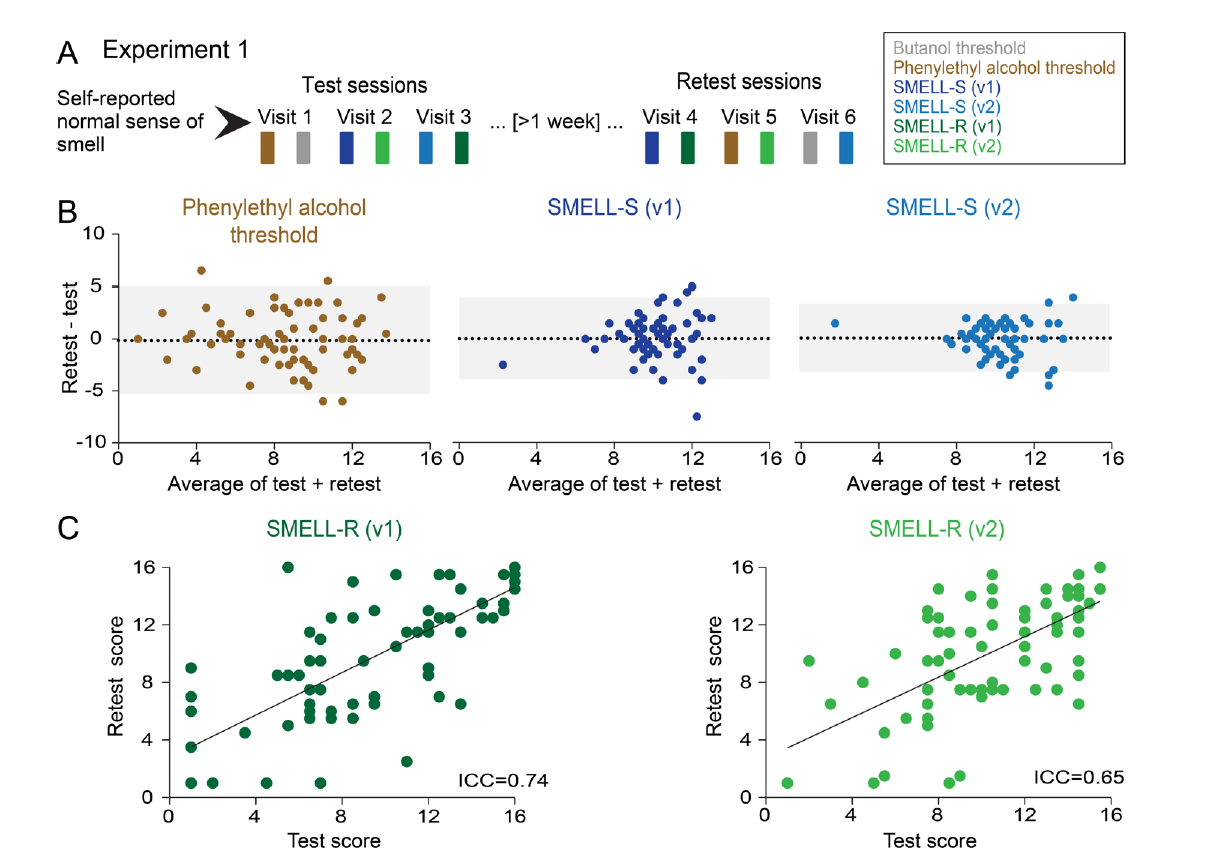
Test-retest reliability and relationship between SMELL-S and SMELL-R tests. (*A*) Experiment 1 design, showing one example of the many different presentations of the six smell tests. SMELL-R tests were always administered after SMELL-S or threshold tests in a given visit. (*B*) Bland-Altman plots of the indicated tests where each dot represents data from one subject. Black dotted lines represent the average of differences between retest and test scores, and gray areas indicate 95% limits of agreement (average difference ± 1.96 S.D. of the difference, n=74-75). (*C*) Test and retest scores for SMELL-R (v1) and SMELL-R (v2) where each dot represents data from one subject (ICC = intraclass correlation coefficient; n=73-75).

### Test-retest reliability

Effective diagnostic tests must be designed with high test-retest reliability. We therefore measured the reliability of SMELL-S and SMELL-R in a population of subjects with a self-reported normal sense of smell (Experiment 1; Fig. *2A*). We tested two versions of SMELL-S and SMELL-R (v1 and v2), which differed in the 30 components used for the mixtures. We also carried out conventional threshold tests with the monomolecular odorants phenylethyl alcohol and butanol. All tests were self-administered with stimuli presented in glass jars. We excluded data from the butanol threshold test from analysis because the stimuli were not stable, as evidenced by a decline in average daily score for all subjects over the course of the four-month study (Supporting Data Set 1).

To assess test-retest reliability for SMELL-S, we computed the absolute difference in test-retest scores for each subject (Fig. *2B*). The bias, as defined by the difference between the average of the test and retest scores, was close to zero for all three tests. This indicates that subjects did not show systematically different performance between the test and retest sessions. The 95% limits of agreement were much smaller for the two SMELL- S tests than the phenylethyl alcohol threshold test (Fig. *2B*). We did not calculate test-retest correlations because of the large inter-individual variability of the phenylethyl alcohol threshold test compared to the two versions of SMELL-S (24). These results demonstrate that the SMELL-S test is more reliable than the phenylethyl alcohol threshold test. The phenylethyl alcohol threshold test is commercially available as Sniffin’ Sticks, a well-validated test administered by clinical staff that utilizes felt-tip pens for odorant delivery (23, 25). To confirm that our self-administered phenylethyl alcohol threshold test presented in glass vials produced results comparable to Sniffin’ Sticks, we re-invited 23 subjects from Experiment 1 and administered the Sniffin Sticks’ version of the phenylethyl alcohol threshold test. There was a strong correlation between the phenylethyl alcohol threshold self-administered in glass vials and Sniffin’ Sticks administered by a research assistant (r=0.87; 95% confidence interval: 0.72 - 0.95, Pearson correlation). This further confirms our conclusions that both versions of SMELL-S are more reliable tests of olfactory sensitivity than thresholds measured with phenylethyl alcohol.

**Fig. 3.**
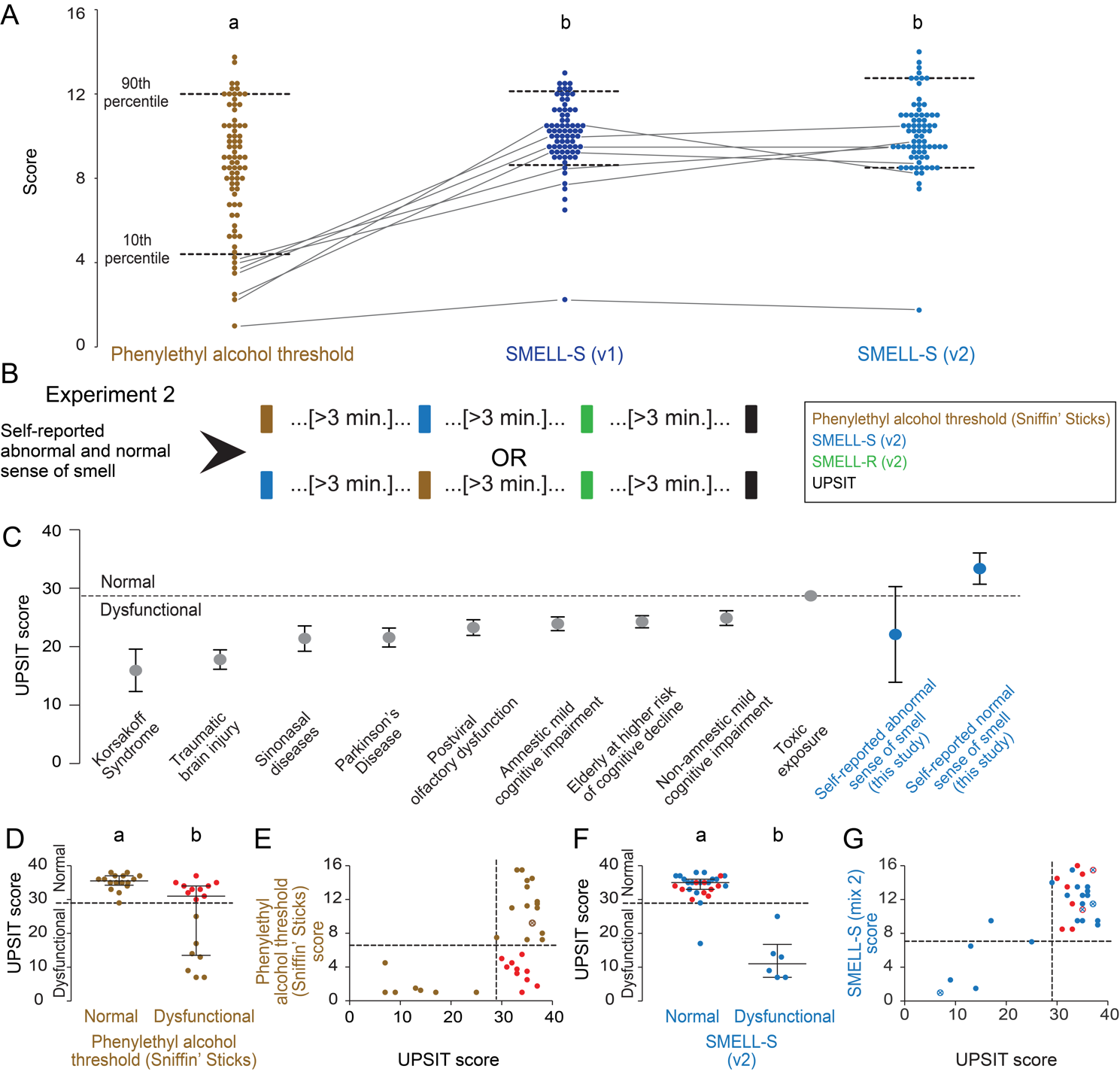
Addressing the problem of odor-specific insensitivity. (*A*) Subject average scores of the indicated tests from Experiment 1 between test and retest. Data marked with different letters indicate significantly different inter-individual variance between groups (p<0.0001; Conover squared ranks test, n=74-75). Data from the seven lowest-scoring subjects in the phenylethyl alcohol test are connected by lines. (*B*) Experiment 2 design. (*C*) The relationship between different etiologies of olfactory dysfunction and UPSIT scores in published studies, as well as of Experiment 2 subjects divided by self-reported smell abilities (mean ± 95% confidence interval). References: Korsakoff Syndrome (43), traumatic brain injury (45), sinonasal disease (45), Parkinson’s Disease (46), postviral olfactory dysfunction (45), amnestic mild cognitive impairment (9), elderly at higher risk of cognitive decline (9), non-amnestic mild cognitive impaired (9), and toxic exposure (47). (*D*) UPSIT scores for subjects divided into normal and dysfunctional groups according to phenylethyl alcohol threshold (Sniffin’ Sticks) (cut-off score= 6.5) performance, where each dot represents data from one subject. Subjects scored as normal by the UPSIT but dysfunctional by Sniffin’ Sticks phenylethyl alcohol test are colored in red in D and E. Data from the remaining subjects are colored brown here and in E. Data labeled with different letters are significantly different (p=0.0003, Mann-Whitney test). Medians and interquartile range are represented. (*E*) Comparison of UPSIT and phenylethyl alcohol (Sniffin’ Sticks) scores. (F) UPSIT scores for subjects divided into normal and dysfunctional groups according SMELL-S (v2) (cut-off score= 7) performance, where each dot represents data from one subject. Subjects scored as normal by the UPSIT but dysfunctional by Sniffin’ Sticks phenylethyl alcohol test in D are colored red in F and G, while the others are in blue. Data labeled with different letters are significantly different (p<0.0001; Mann-Whitney test), and represented as median and interquartile range. (G) Comparison between UPSIT and SMELL-S (v2) scores. Subjects with identical values are indicated by superimposed open circles and an X, and retain the specified color coding.

We next examined the test-retest reliability of the SMELL-R test. Because the interindividual variability between SMELL-R (v1) (mean 9.3 ± 4.3 standard deviation) and SMELL-R (v2) (mean 10.2 ± 3.5 standard deviation) did not differ significantly (p= 0.074, F test) (Fig. *2C*), we calculated the intraclass correlation coefficient (ICC) for the SMELL-R tests. By this metric the two versions of SMELL-R are very reliable (Fig. *2C*). Because SMELL-R (v2) had lower interindividual variability, we used this version of the test for the remaining experiments in the study.

### Addressing the problem of odor-specific insensitivity

Smell tests that use a monomolecular stimulus like phenylethyl alcohol may misdiagnose a subject with odor-specific insensitivity to this odorant. A test based on mixtures of many components would overcome this problem, because each odorant in the mixture has only a small impact on the overall test score (26). Such a test would be highly effective in diagnosing general olfactory dysfunction, rather than variability in sensitivity to any individual odorant.

To explore how odor-specific sensitivity affects the accuracy of smell dysfunction diagnosis, we compared the performance of subjects in Experiment 1 on smell tests that used monomolecular stimuli or mixtures. The variability in test scores across all subjects in Experiment 1 of the phenylethyl alcohol threshold was significantly higher than that of SMELL-S (v1) and SMELL-S (v2) (Fig. *3A*). Of the 7 subjects in the lowest 10 ^th^ percentile in the phenylethyl alcohol threshold test, only one was in the lowest 10 ^th^ percentile for both versions of SMELL-S. We speculate that this subject has an impaired sense of smell. The low phenylethyl alcohol scores of the other 6 subjects likely reflect odor-specific insensitivity to odorant rather than impaired olfactory function. These subjects would have been misdiagnosed with smell dysfunction using the phenylethyl alcohol threshold test.

To confirm this, we compared the SMELL-S test with the Sniffin’ Sticks phenylethyl alcohol threshold test and the North American version of the UPSIT (Experiment 2; Fig. *3B*). Since SMELL-S (v2) had the narrowest 95% limits of agreement (Fig. *2B*), we used this version of SMELL-S for the rest of this study. In Experiment 2, we assessed the performance of subjects with a self-reported normal or abnormal sense of smell on the UPSIT, Sniffin’ Sticks, and SMELL-S (v2). Based on results in Figure *3A*, we anticipated that SMELL-S (v2) would be more accurate than the Sniffin’ Sticks phenylethyl alcohol threshold test in identifying subjects with smell dysfunction. We used the UPSIT to benchmark the performance of the Sniffin’ Sticks phenylethyl alcohol threshold test compared to SMELL-S (v2). Because this smell test is composed of 40 different items, the final score is not strongly affected by odor-specific insensitivity to any given stimulus among the 40 items of the test. To set a cut-off between normal and dysfunctional subjects, we performed a literature search on mean UPSIT scores in North American patients suffering from smell dysfunction caused by various etiologies. Based on this analysis and consistent with an earlier study (3), we defined normal olfactory function as an UPSIT score of 29 and over and smell dysfunction as an UPSIT score of 28 and lower (Fig. *3C*). In Experiment 2, the mean score of the subjects with self-reported smell dysfunction was below this cut-off, whereas the mean score of those with self-reported normal sense of smell was above the cut-off (Fig. *3C*). For the Sniffin’ Sticks phenylethyl alcohol threshold test we used the cut- off specified by the manufacturer, with normal defined as a score higher than 6.5 and dysfunctional a score of lower than 6.5.

Subjects in Experiment 2 were divided into normal and dysfunctional according to their performance on the Sniffin’ Sticks phenylethyl alcohol threshold test (Fig. *3D*). Using the UPSIT cut-off score as a metric of olfactory dysfunction, 10 subjects would have been misdiagnosed as having olfactory dysfunction by the Sniffin’ Sticks phenylethyl alcohol threshold test (Fig. *3D-G*, red dots). When we divided subjects according to performance on the SMELL-S (v2) test using a cut-off score of 7 (see Fig. *4*), we found that only one subject was given a different diagnosis using the UPSIT than with SMELL-S (v2) (Fig. *3F*). This demonstrates that the SMELL-S (v2) test is superior to the Sniffin’ Sticks phenylethyl alcohol threshold test in accurately diagnosing smell dysfunction. We conclude that odor-specific insensitivity to phenylethyl alcohol makes the Sniffin’ Sticks phenylethyl alcohol threshold unreliable, and that the use of odor mixtures in SMELL-S (v2) overcomes this problem and is superior in accurately diagnosing smell dysfunction.

### Diagnostic accuracy

To be useful in the clinic, smell tests must correctly identify patients with smell dysfunction, and not misdiagnose normal subjects. In other words, diagnostic tests must balance false positive and false negative results. To establish a diagnostically optimal cut-off score for SMELL-S (v2) and SMELL- R (v2), we divided subjects into dysfunctional and normal using an UPSIT cut-off score of 29, and examined SMELL-S (v2) scores of self-reported normal and abnormal subjects in these two groups (Fig. *4A*). Subjects with normal UPSIT scores had significantly higher SMELL-S (v2) scores than those that were dysfunctional (Fig. *4A*). Subjects with a self-reported normal sense of smell (blue dots in Fig. *4A*) had significantly higher SMELL-S (v2) scores (median, 12.5; interquartile range, 11-14) than whose with self-reported abnormal sense of smell (median, 7.75; interquartile range, 2.25- 9.50) (red dots in Fig. *4A*) (p=0.0011, Mann-Whitney test).

We next determined the overall accuracy of SMELL-S (v2), and selected an optimal cut-off score to differentiate normal and dysfunctional subjects (Fig. *4B-C*). The standard measure of clinical test accuracy is the area under the receiver operating characteristic (ROC) curve, which plots the true and false positive rates at different cut-off scores. The area under the ROC curve of SMELL-S (v2) is 0.98 (95% confidence interval: 0.85-1.00) (Fig. *4B*), which is close to the perfect accuracy of 1.

To select the cut-off value for SMELL-S (v2) that optimally distinguishes normal and dysfunctional subjects we calculated Youden’s Index (27) at each of 14 SMELL-S (v2) cut-off scores. A Youden’s Index value of 1 indicates no false positives and no false negatives. (Fig. *4C*). Based on this analysis, the administration of SMELL-S (v2) with a cut-off value of 7 will be clinically useful for physicians to diagnose patients with olfactory dysfunction. We carried out the same procedure to determine the accuracy of the SMELL-R olfactory resolution test. Subjects classified as dysfunctional by their UPSIT score had lower SMELL-R (v2) scores, and subjects classified as normal by UPSIT performance had a higher SMELL-R (v2) scores (Fig. *4D*). The area under the ROC curve for SMELL-R (v2) was 0.82 (95% confidence interval: 0.65- 0.93) (Fig. *4E*). The optimal cut-off assessed by Youden’s Index was 8.5 (Fig. *4F*). These data show that SMELL-R at a cut-off value of 8.5 will be clinically useful for diagnosing smell dysfunction.

### Addressing the problem of different prior olfactory experiences

A major goal of this study was to develop a test that does not have to be adapted to different populations. To ask if SMELL-R (v2) performs well in different countries, we compared SMELL-R (v2) performance between Taiwanese and North American subjects (Experiment 3, Fig. *5A*). As a positive control we used the North American version of the UPSIT for both populations, because previous work has shown that Taiwanese subjects have systematically lower scores on this test due to unfamiliarity with several of the test items (18). To enable self-administration of the UPSIT we supplied Taiwanese subjects with a Chinese translation of the English multiple-choice questions in the test booklet. SMELL-R (v2) did not require any language translation because it is non-semantic.

As expected, North Americans performed better on most of the items in the UPSIT, with the biggest differences found for “pine”, “lime”, “cherry”, and “rose” (Fig. *5B*). Even so, several items were frequently mistaken by North American subjects, including “paint thinner” when the correct answer was “cheddar cheese”, “musk” instead of “lime”, and “wintergreen” instead of “bubble gum”. The Taiwanese subjects also struggled with the “cheddar cheese” item, also frequently mistaking it for “paint thinner”, but in addition mistook “turpentine” for “soap”, “motor oil” for “grass”, and “clove” for “licorice”. The overall UPSIT scores for Taiwanese subjects were significantly lower than those of the North American subjects (Fig. *5C*) (p<0.0001, Mann Whitney test). In contrast, Taiwanese subjects scored higher on SMELL-R (v2) than the North American subjects (Fig. *5D*) (p=0.0157, Mann Whitney test). The difference for the two populations was much smaller for SMELL- R (v2) than the UPSIT as determined by calculating the difference in Z-scores (Fig. *5E*). While we do not know the underlying cause for the superior performance of Taiwanese subjects on SMELL-R (v2), the results show that our test avoids the bias seen for the UPSIT, in which test performance is systematically higher in the population for which the test was developed. We conclude that SMELL-R (v2) can be applied across different populations with different prior olfactory experiences, and without the need to adapt it to the local culture and language.

**Fig. 4.**
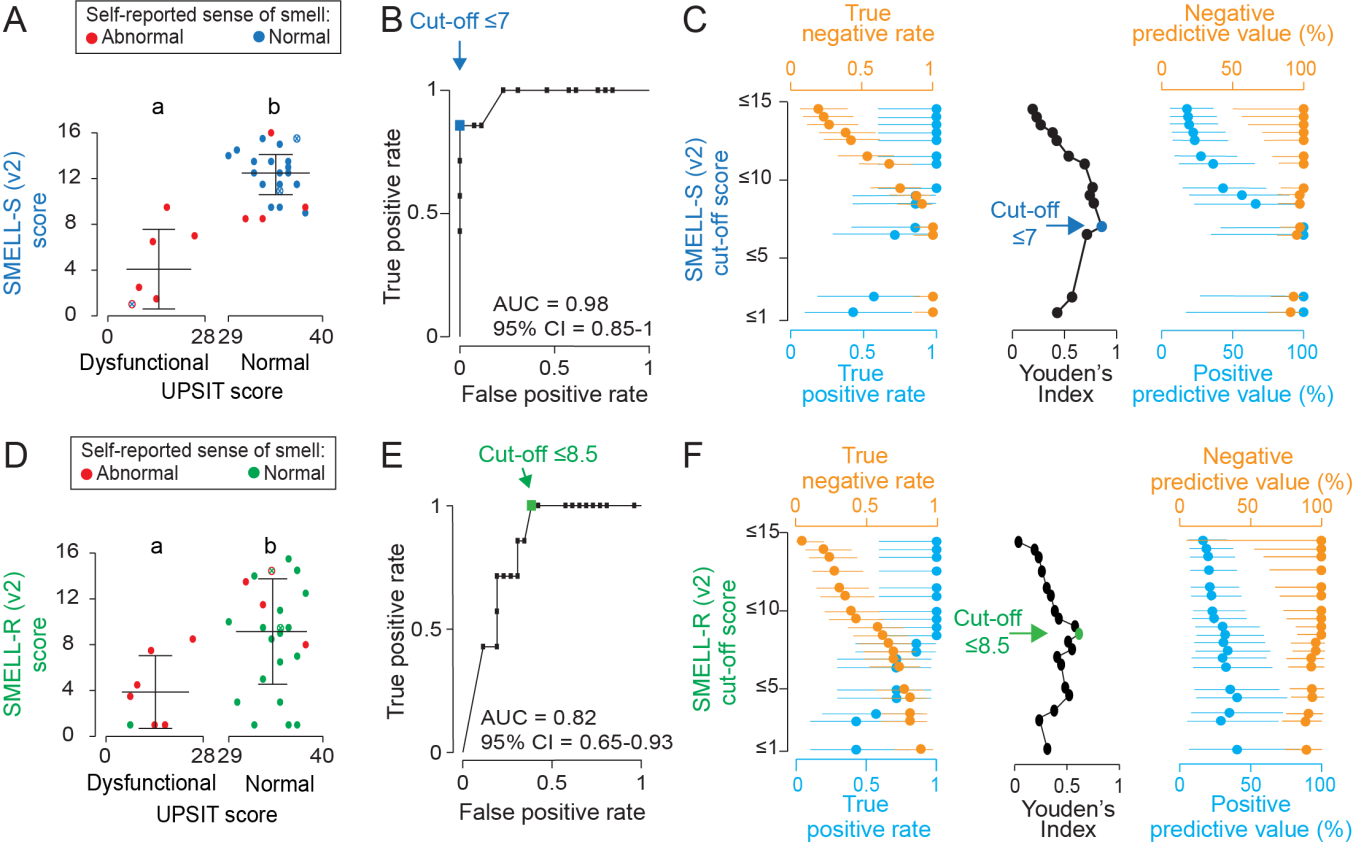
SMELL-S and SMELL-R diagnostic accuracy. (*A*) Comparison of UPSIT and SMELL-S (v2) scores for Experiment 2 subjects (mean ± S.D.) Subjects were divided into dysfunctional (n=7) and normal (n=26) using an UPSIT cut-off score of 29. Data labeled with different letters are significantly different (p=0.0005, two-sided unpaired t-test with Welch’s correction). (*B*) Area under the ROC curve (AUC) for SMELL-S (v2). The optimal cut-off is indicated by the blue dot. (*C*) Plots of four measures of diagnostic accuracy resulting from different cut-off scores for SMELL-S (v2) (percentage ± 95% confidence interval). The optimal cut-off score for olfactory dysfunction defined by Youden’s Index (center) is indicated by the blue dot. (*D*) Comparison of UPSIT and SMELL-R (v2) scores for Experiment 2 subjects (mean ± S.D.) Subjects were divided into dysfunctional (n=7) and normal (n=26) using an UPSIT cut-off score of 29. Data labeled with different letters are significantly different (p=0.0035, two-sided unpaired t-test with Welch’s correction). Subject Expt 2- A006 had a self-reported normal sense of smell but low scores on the UPSIT, as well as SMELL-S (v2) in *A* and SMELL- R (v2) in *D*. (*E*) Area under the ROC curve (AUC) for SMELL-R (v2). The optimal cut-off is indicated by the green dot. (*F*) Plots of four measures of diagnostic accuracy resulting from different cut-off scores for SMELL-R (v2) (rate or percentage ± 95% confidence interval). The optimal cut-off score for olfactory dysfunction defined by Youden’s Index (center) is indicated by the green dot. Subjects with identical values are indicated by superimposed open circles and an X, and retain the specified color coding.

**Fig. 5.**
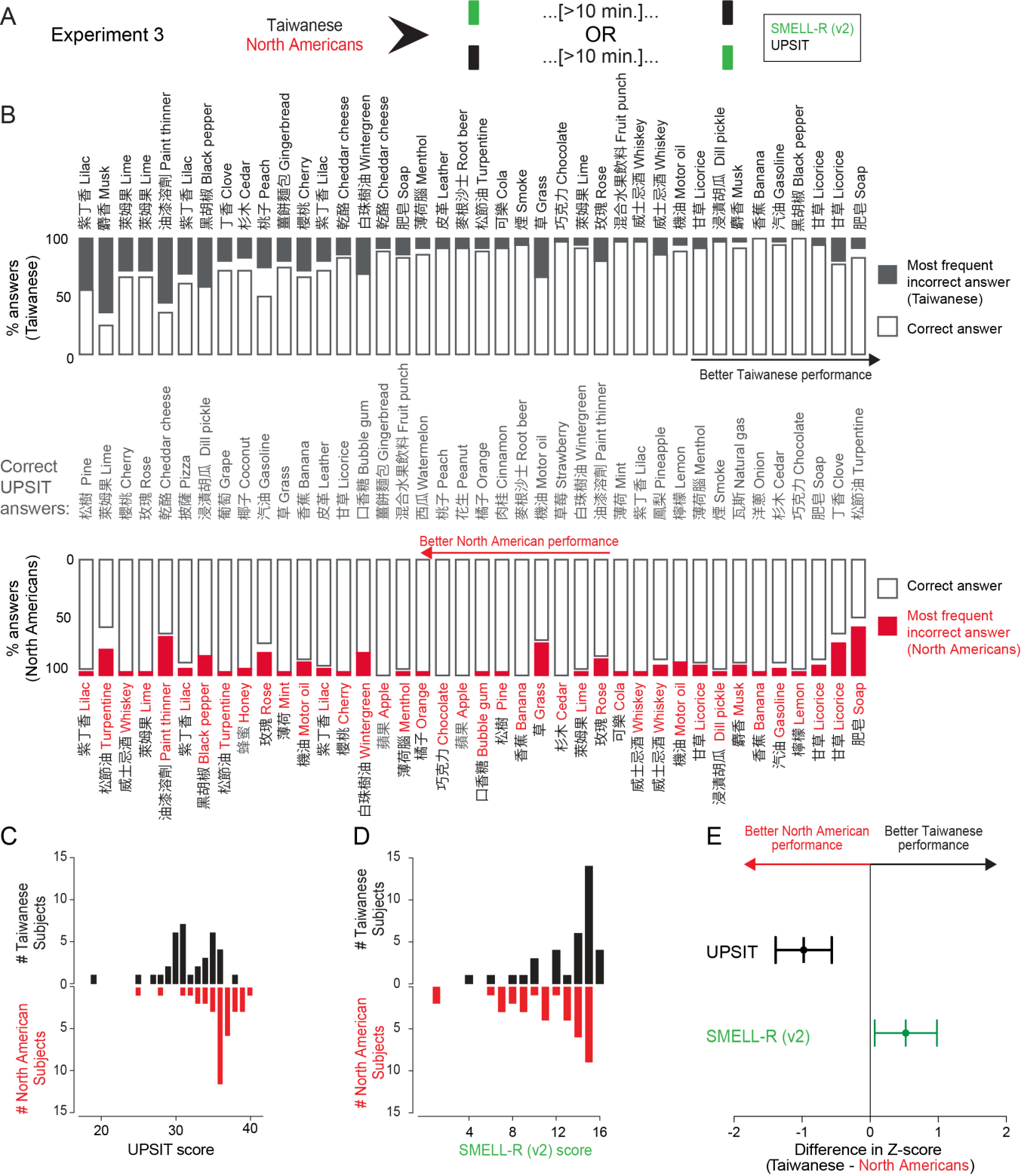
Addressing the problem of different prior olfactory experiences. (*A*) Experiment 3 design. (*B*) Performance of Taiwanese (n=36) (top) and North American (n=36) (bottom) subjects for individual UPSIT items, open bar bars indicate correct answers and solid bars indicate the most frequent incorrect answer. (*C-D*) Histogram of North American and Taiwanese subject scores for the UPSIT (C) and SMELL-R (v2) (D). (*E*) Cross-population comparison of UPSIT and SMELL-R (v2) (mean ± 95% confidence interval) for subjects in (C,*D*).

## Discussion

In this study, we addressed current limitations in clinical testing for olfactory dysfunction by developing effective smell tests that do not misdiagnose subjects with odor-selective insensitivity, and that can be utilized with diverse populations across the world.

The first objective of this work was to eliminate the problems inherent in olfactory sensitivity tests that rely on a single molecule. Although it is well known that specific insensitivity to individual odorant molecules is common in normal human subjects (28), commercial threshold tests use mono- molecular stimuli such as butanol or phenylethyl alcohol to test olfactory sensitivity (23, 29). Our data suggest that this approach confounds specific and general olfactory sensitivity. Previous studies showed a lack of correlation between butanol and phenylethyl alcohol threshold, further demonstrating that current commercial threshold tests measure odor-specific sensitivity rather than general olfactory function (17). We show here that the solution to this problem is to use mixtures of molecules instead of single molecules. We and other authors have shown that the inter- and intra-individual variability in threshold scores was reduced, and test-retest reliability was increased by testing olfactory sensitivity with odor mixtures rather than single molecules (26, 30). SMELL-S is a reliable, accurate, and effective method for measuring olfactory function without conflating general loss of smell sensitivity and specific insensitivity to an odorant.

A second objective of this project was to introduce a test of olfactory resolution (SMELL-R) that quantifies olfactory discrimination ability. Auditory and visual stimuli used in the clinic differ by tone frequency or letter size, leading to quantitative and standardized diagnostic tests such as the audiogram and the eye chart. In olfaction, it is more complicated to quantify similarity between olfactory stimuli. Currently available discrimination tests consist of several pairs of odorants that must be discriminated by the patient. There currently is no method to quantify how difficult the individual discrimination tests are. Is distinguishing “rose” from “leather” more or less difficult than discriminating “pineapple” from “licorice”? To overcome this problem, we used a physical scale based on the number of shared components between two mixtures. The more components two mixtures share, the more difficult it is to discriminate them (21, 22). By using this physical scale, we reliably determined the resolution of the olfactory system.

A third objective was to develop smell tests that utilize stimuli that have not been previously encountered by patients to minimize the influence of prior olfactory experience on the test results (19, 20). We accomplished this by using mixtures of 30 different molecules. These exact mixtures are very unlikely to be encountered outside the laboratory. Furthermore, before mixing them, the chemicals were diluted so that they had approximately equal intensity to ensure that the percept of the mixture is not dominated by a single odorant. The resulting smell of such mixtures has been described as olfactory white (21). Using these stimuli is an improvement over the use of odorants that can be readily linked to their usual source, but only by those who have prior experience with it (18, 31, 32).

Developing an olfactory test to reliably diagnose smell dysfunction is of great clinical importance not only because of the negative effects of smell dysfunction on quality of life, but because olfactory dysfunction may be an effective biomarker for predicting Alzheimer’s disease (9). Piriform and entorhinal cortex, brain regions affected in the early stages of Alzheimer’s disease (33–35), are also involved in the processing of olfactory information. Consistent with this anatomical observation, Alzheimer’s disease patients show both reduced sensitivity to test odorants and a reduced ability to match specific odorants with words describing them (36). We suggest that the non-semantic SMELL-R and SMELL-S test will be an effective and non-invasive method to identify Alzheimer’s disease in patients before the earliest symptoms of memory loss are detected.

## Acknowledgements

We thank our research volunteers for their time and interest in the study and the staff of The Rockefeller University Hospital Outpatient Clinic for invaluable support. Chris Vancil provided custom programming for the Rockefeller University Smell Study smell test computer interface, and Joel M. Correa da Rosa and Caroline Jiang provided expert biostatistical guidance. Yuanbo Wang provided a script to compute test scores in Experiment 1. We thank Barry Coller, Ashutosh Kacker, Kevin Lee, and members of the Vosshall Lab for discussion and comments on the manuscript. This work was funded by the National Center for Advancing Translational Sciences (NCATS), National Institutes of Health (NIH) Clinical and Translational Science Award (CTSA) program UL1 TR000043. A.K. is the recipient of a Branco Weiss Science in Society Fellowship. L.B.V. is an investigator of the Howard Hughes Medical Institute.

## Materials and Methods

### General, subjects

All behavioral testing with human subjects took place between March 2015 and December 2016, and was approved and monitored by the Institutional Review Boards of The Rockefeller University in New York, NY (USA), except the Taiwanese arm of Experiment 3, which was approved by the Institutional Review Board of Taichung Veterans General Hospital in Taichung, Taiwan. North American subjects were recruited by The Rockefeller University Clinical Research Recruitment and Outreach Support Service (37). Taiwanese subjects were recruited by the nursing staff of the Department of Otorhinolaryngology at the Taichung Veterans General Hospital (Taiwan). All subjects gave their written informed consent to participate in these experiments, and were compensated for their time. All North American and Taiwanese subjects were able to understand and follow instructions in English or Mandarin, respectively. Subjects were aged 18 or over and agreed to refrain from using perfume or cologne, and ingesting anything except water one hour prior to the study visit. At the beginning of each visit, subjects washed their hands with odorless soap. For subjects reporting a normal sense of smell and taste, we excluded subjects who presented with current or past history of conditions that might be related to smell loss (acute or chronic rhinosinusitis, nasal tumor, upper respiratory tract infection or head trauma that altered the sense of smell for more than one month, history of brain or sinonasal surgery, asthma, stroke, neurodegenerative disease, radiation therapy or chemotherapy, active smoking, or consumption of medication affecting the sense of smell during the study). Participants with self-reported smell dysfunction were not subject to these exclusion criteria. All raw data in the paper, including details about the demographics of the subjects, odorants, and composition of the test stimuli are in Supporting Data Set 1.

### General, tests

To allow for self-administration and automatic data collection, we designed a custom computer application that was used for the vial threshold tests and also the SMELL-S and SMELL-R tests. The testing station comprised a computer, wireless mouse, barcode scanner, and trays containing numbered stimulus containers labeled with bar codes. Triangle tests were set up so that subjects were never tested with the same set of stimuli twice in a row, to avoid the situation where subjects remembered their answers from the previous trial. Subjects used a barcode scanner to register test data automatically. Subjects took between 20-35 minutes to complete each smell test, with the exception of the UPSIT which took 10-15 minutes. A standard inter-trial interval was imposed to avoid odor adaptation by requiring subjects to play a computer game for 20 seconds.

SMELL-S and SMELL-R were created with four different mixtures of 30 molecules drawn from a panel of 109 monomolecular, intensity-matched chemicals. These odorants were selected from stimuli utilized in previous psychophysical studies (21, 38). We used only molecules that minimally activated the trigeminal system, because such stimuli can be detected by anosmic subjects (39, 40). A characteristic of trigeminal activation by a molecule is a fresh, cold, burning, eucalyptus, pungent, or tickling sensation. We used a lateralization task in which an odorant is applied into only one nostril to assign a lateralization score to each molecule. It is possible to localize the stimulated nostril if it activates the trigeminal system. In contrast, it is much harder to localize an olfactory stimulus (41). Lateralization tasks were self-administered by one investigator. Two disposable squeeze bottles were placed in a device facilitating simultaneous squeezing and stimuli delivery in each nostril. Only one bottle was filled with an odor stimulus. The tip of each bottle was fitted with a foam piece that conformed to the investigator’s nostril, and placed at the entrance of each nostril. The investigator squeezed both bottles simultaneously and attempted to localize which nostril had received the stimulus. After each task, the device was spun on a rotating platform to randomize the odor-stimulus side. The final score corresponded to the number of correct tasks. There were a total of 20 tasks (42). As a control experiment, we found that the lateralization score of the trigeminal stimulus eucalyptol (CID: 2758) at pure concentration was high (median, 20; interquartile range, 19.25 – 20; 4 trials). The lateralization score of the olfactory stimulus vanillin (CID: 1183) at pure concentration was low (median, 6.5; interquartile range, 5-12.5, 6 trials). The difference between the lateralization scores of eucalyptol and vanillin was statistically significant (p=0.0009, Mann-Whitney test). Each candidate for the mixtures was tested once. We included candidates with a score of 11 and below in the design of the mixtures (Supporting Data Set 1).

To intensity-match molecules to be used in mixtures, odorants were diluted and three investigators individually classified them as too weak, well matched, or too strong. The concentration of “too weak” stimuli was increased and that of “too strong” stimuli decreased by a factor of 10. Weak components that could not be intensity-matched even at pure concentrations were excluded from the pool of odorants. We repeated this process until most of the components fell into the optimal intensity range. For 18 components investigators could not reach a consensus about intensity, but these were nevertheless used in the mixtures (CID: 1068, 7969, 31244, 9589, 17898, 104721, 3314, 14491, 62144, 7583, 7983, 60999, 251531, 7799, 61151, 9609, 8118, 89440). With these components, we created four mixtures of 30 components. The SMELL-S (v1) mixture was used as the ODD odor in SMELL-R (v1), and the SMELL-S (v2) mixtures was used as the CONTROL odor in SMELL R (v2). The mixtures for SMELL-R (v1) CONTROL odor and SMELL-R (v2) ODD odor were unique to these tests. Details of all mixtures are in Supporting Data Set 1.

Stimuli for the vial threshold tests and SMELL-S were presented to subjects with amber glass vials (height: 95 mm, diameter: 28 mm). Stimuli for SMELL-R were presented to subjects with amber glass jars (height: 51 mm, diameter: 55 mm). The complete list of stimuli used in this study is in Supporting Data Set 1.

### Threshold tests: phenylethyl alcohol and butanol

Threshold tests were administered as a series of triangle tests. Subjects were presented with three vials: two contained 1 ml solvent (paraffin oil) and one contained either phenylethyl alcohol or butanol diluted in solvent in a total volume of 1 ml. Tests comprised 16 different concentrations generated by serial dilutions (1:2) of either odorant in paraffin oil, with the starting concentrations at 0.0313% for phenylethyl alcohol (vial) and 0.25% for butanol (vial). The subject was prompted to sniff each vial and select the one with the strongest perceived odor using an adaptive staircase procedure commonly used in smell testing (23). If they were unable to detect any difference among the three vials, they were prompted to choose one at random. The procedure started at the lowest concentration. If they identified an incorrect vial, the second next higher concentration was presented and so on, until they identified the correct vial. If the subjects identified the correct vial, they were retested at the same concentration. If they identified the correct vial in this retest, they were tested at the next lower concentration. If they identified an incorrect vial, they were tested at the next higher concentration. A reversal is when the direction in which the concentration is changed reverses. The procedure ended after the seventh reversal, or until the subject failed the level with the highest concentration twice in row, or succeeded with the lowest concentration level 5 times in row. The threshold was defined as the average of the concentrations at which the last two reversals occurred. If the highest concentration were not correctly identified twice, the score was 1. If the lowest was identified 5 times in a row, the score was 16.

### SMELL-S olfactory sensitivity test (v1 and v2)

For SMELL-S (v1) and SMELL-S (v2), we prepared 19 serial dilutions in paraffin oil (1:2) of two different mixtures of 30 monomolecular odorants and used the last 16 dilutions, such that the tests ranged from easiest (level 1, 1:8 dilution) to most difficult (level 16, 1:262,144 dilution). Subjects were asked to sniff 3 vials, one of which was filled with 1 ml of a mixture of 30 components, and the other two were filled with 1 ml of solvent (paraffin oil). Subjects were instructed to pick out the one vial with the strongest perceived odor. If they were unable to detect any difference among the three vials, they were prompted to choose one at random. The procedure started at the lowest concentration (level 16). We calculated the SMELL-S sensitivity score following the same adaptive staircase procedure described above. For each subject, we measured the olfactory sensitivity with two versions of the tests, SMELL-S (v1) and SMELL-S (v2), which differed only by the chemical composition of the mixtures.

### SMELL-R Olfactory Resolution tests (v1 and v2)

For SMELL-R (v1) and SMELL-R (v2), we prepared 16 pairs of mixtures of 30 monomolecular odorants that differ in how many components the two mixtures in the pair share from 0% (easiest; level 1) to 96.7% (most difficult; level 16). To create 16 levels of increasing overlapping components, we randomly progressively replaced components of a mixture of 30 molecules (we termed this the ODD odor), with components from another mixture of 30 components that did not change in composition across the levels (we termed this the CONTROL odor). Increasing the level of difficulty by one point correspond to an addition of 2 overlapping molecules between both mixtures, except from level 15 to 16, where we added only 1 shared molecule. Stimuli (8 ml) were introduced into jars containing absorbent cotton pads. Subjects were asked to sniff the contents of 3 jars, one of which was filled with 8 ml of a mixture of 30 components, and the other two were filled with 8 ml of a mixture of 30 components with different degree of overlap with the first jar. Subjects were instructed to pick out the odd jar. If they were unable to detect any difference among the three jars, they were prompted to choose one at random. Triangle tests started at a medium difficulty (level 8). If they identified the incorrect jar, the next easier level was presented. We calculated the SMELL-R resolution score following the same adaptive staircase procedure described above. For each subject, we measured the olfactory resolution with two versions of the tests, SMELL-R (v1) and SMELL-R (v2), which differed only in the chemical constituents of the two sets of mixtures.

### Sniffin’ Sticks phenylethyl alcohol threshold test

The Sniffin’ Sticks (23) threshold phenylethyl alcohol threshold test is a commercial product that uses felt-tip pens filled with odorant instead of ink for odor presentation. [threshold module (2-phenyl ethanol) of the extended Burghart Sniffin’ Sticks test, Burghart Messtechnik, item # LA-13-00015]. The test comprises pens containing 16 serial dilutions of phenylethyl alcohol (1:2) in solvent (propylene glycol) with a starting concentration of 4%. The test was administered as a triangle test. Three pens were presented to the subjects by the investigator in a randomized order. Two pens contained the solvent only, and the third pen contained the diluted odorant. Subjects were blindfolded with a disposable mask because the color code of the Sniffin’ Sticks reveals which pen contains the odor, and were asked to identify the pen with the strongest perceived odor. The procedure started at the lowest or second lowest concentration of odorant (level 16 or 15, respectively). We calculated the threshold score following the same adaptive staircase procedure described above except that the threshold was defined as the average of the last four reversals occurred.

### UPSIT

The University of Pennsylvania Smell Identification Test (UPSIT, marketed as the Smell Identification Test by Sensonics International) is a well-validated and self-administered smell identification test widely used in the USA (43). The test consists of 4 different 10-page booklets, with a total of 40 monomolecular stimuli. On each page, there is a different scratch and sniff strip which is embedded with a microencapsulated odorant. There are also four choice multiple-choice questions on each page. Subjects used the tip of a pencil to release the smell of the stimuli. Subjects sniffed the odorant and selected the one word among the four options (for example “paint thinner”, “cherry”, “coconut”, or “cheddar cheese”) that most closely matched their perception of the smell. Subjects entered their answers to the 40 multiple-choice questions manually into a booklet, and investigators transferred the data manually into a spreadsheet. UPSIT performance was scored as the number of correct answers out of 40. We used the same North American UPSIT (43) on subjects at Rockefeller University and Taichung Veterans General Hospital. The Taiwanese subjects were given a reference sheet on which the English multiple-choice questions in the UPSIT booklets were translated into Chinese by R.-S. J. (Fig. *5B*) (18).

### Experiment 1, design

In this protocol (Rockefeller University IRB Protocol # JHS-0862), we studied the test-retest reliability of SMELL-S and SMELL-R. We invited volunteers with self-reported normal sense of smell and taste to the Rockefeller University Hospital for six visits (Fig. *2A*). During these six visits, six olfactory tests were performed, each of them once during a test session (visit 1 to 3), and then again during a retest session (visit 4 to 6). There was a gap of at least 1 week between the last test visit (visit 3) and the first retest visit (visit 4), and a gap of at least 24 hours between each of the other visits. At each visit, two of the six tests were performed. Although the order of the tests was randomized, in any visit where SMELL-R tests were administered, they were always administered after the SMELL-S or the threshold tests. This experiment was done between March and June 2015.

### Experiment 1, subjects

75 subjects (43 female) participated in this experiment, with a mean age of 44 (range: 21-74). 34 subjects self-identified as White, 26 as Black, 6 as Asian, 2 as Mixed Race, and 7 as Other. 11 subjects self-identified as Hispanic. It took an average of 21 days (range: 14-38 days) for subjects to complete all 6 visits in this experiment.

### Experiment 1, statistical analysis

The Intra-class Correlation Coefficient (ICC) was used to measure absolute agreement between test and retest measures for the whole cohort. A sample of n=75 subjects provided 95% confidence that the ICC in the population was larger than 0.67 based on a sample distribution that is centered on 0.8 (44). Bland-Altman plots were used as an auxiliary tool if significant differences in inter-individual variability were found between compared tests (24) (Fig. *2B*). We used the non-parametric Conover squared ranks test to assess equality of variance across threshold tests. Statistical significance was reached when p<0.05 (Fig. *3A*).

### Experiment 2, design

This experiment was carried out under Rockefeller University IRB protocol #JHS-0922, and was designed to evaluate the accuracy of our tests and whether SMELL-S can distinguish between subjects with specific-anosmia to phenylethyl alcohol but an otherwise normal sense of smell and subjects with smell dysfunction. During a single visit in December 2016, subjects performed four smell tests. The first two tests were either SMELL-S (v2) or Sniffin’ Sticks phenylethyl alcohol threshold test. The order of these first two tests was randomized. It was followed by SMELL-R (v2) and finally the UPSIT, as a validated commercial reference test. The investigators enforced a break of at least 3 minutes between tests. During some of the breaks, participants filled out a questionnaire to provide demographic information and answer questions about their sense of taste and smell (Supporting Data Set 1). In 7 cases in the UPSIT tests in Experiment 2, subjects did not provide an answer to a given item, and this was scored as an incorrect answer. The missing data correspond to 3 subjects who missed one item each, and 2 subjects who missed 2 items each.

### Experiment 2, subjects

This experiment included 33 subjects (22 female), with a mean age of 48 (range: 21-76). 17 subjects self-identified as White, 8 as Black, 3 as Asian, 2 as Mixed Race, 1 as Other. Two subjects opted out of self-reporting race. Four subjects self-identified as Hispanic. We re-enrolled 23 subjects from Experiment 1 who self-reported a normal sense of smell and taste. These 23 were selected based on their threshold test results to have approximately even representation of subjects with low, medium, and high sensitivity to phenylethyl alcohol. In addition, we recruited 10 subjects with self-reported smell dysfunction. The self-reported etiologies are reported in Supporting Data Set 1.

### Experiment 2, statistical analysis

We performed a power analysis and determined that a study with 32 subjects (8 with smell loss and 24 with a normal sense of smell) guarantees 80% power at 5% significance to detect an area under the ROC curve greater than 0.78. Since our actual study included 33 subjects, we carried out a post hoc power analysis using the parameters above to show that we can detect an area under the ROC curve greater than 0.79. We employed Youden’s Index (27) to find the best cut-off score for SMELL- S and SMELL-R to maximize correct classification of the olfactory sensitivity and resolution of a subject, respectively (Fig. *4C,F*). We used two-sided unpaired t-test with Welch’s correction to test for differences between SMELL-S and SMELL-R score in normal and dysfunctional groups (Fig. *4A,C*).

### Experiment 3, design

In this experiment, we investigated how SMELL-R performs on different populations by comparing Taiwanese (IRB TCVGH #CE16119B) and North American (Rockefeller University IRB Protocol #JHS-0901) subjects. The North American subjects were tested at The Rockefeller University Hospital, and the Taiwanese subjects were tested in the Department of Otolaryngology at Taichung Veterans General Hospital. The experimental design was the same in both institutions. Each subject came to the test site for a single visit, during which subjects performed the SMELL-R (v2) and UPSIT, separated by a 10 minute break, in randomized order (Fig. *5A*).

### Experiment 3, subjects

36 subjects were recruited at both sites. All subjects were born and raised in their respective country, had never travelled to the opposite country, and had a self-reported normal sense of smell and taste. In the North American group, the mean age was 25 (range: 19-30), 23 of 36 subjects were female, and 8 self-identified as White, 14 as Black, 4 as Asian, 9 as Mixed Race, and 1 as American Indian or Alaska native. Six self-identified as Hispanic. In the Taiwanese group, the mean age was 26 (range: 19-30) and 26 of 36 subjects were female. Although we recruited subjects with a self-reported normal sense of smell, two of the North American subjects had UPSIT and SMELL-R (v2) scores below the cut-off for olfactory dysfunction (Fig. *5C,D*).

### Experiment 3, statistical analysis

We used unpaired t-test with Welch’s correction to test for differences between smell test performance between North American and Taiwanese subjects (Fig. *5C,D*). These tests were applied with and without the 2 North American subjects with UPSIT scores below the cut-off defined for a normal sense of smell.

### Statistical analysis

Normality of data was tested throughout using the Kolmogorov–Smirnov test, and the appropriate statistics were used according to the distribution of the data. SPSS (IBM) and Prism (Graphpad) was used for all statistical analysis.

